# Feeders and Expellers, Two Types of Animalcules With Outboard Cilia, Have Distinct Surface Interactions

**DOI:** 10.1101/2024.06.29.601328

**Authors:** Praneet Prakash, Marco Vona, Raymond E. Goldstein

## Abstract

Within biological fluid dynamics, it is conventional to distinguish between “puller” and “pusher” microswimmers on the basis of the forward or aft location of the flagella relative to the cell body: typically, bacteria are pushers and algae are pullers. Here we note that since many pullers have “outboard” cilia or flagella displaced laterally from the cell centerline on both sides of the organism, there are two important subclasses whose far-field is that of a stresslet, but whose near field is qualitatively more complex. The ciliary beat creates not only a propulsive force but also swirling flows that can be represented by paired rotlets with two possible senses of rotation, either “feeders” that sweep fluid toward the cell apex, or “expellers” that push fluid away. Experimental studies of the rotifer *Brachionus plicatilis* in combination with earlier work on the green algae *Chlamydomonas reinhardtii* show that the two classes have markedly different interactions with surfaces. When swimming near a surface, expellers such as *C. reinhardtii* scatter from the wall, whereas a feeder like *B. plicatilis* stably attaches. This results in a stochastic “run-and-stick” locomotion, with periods of ballistic motion parallel to the surface interrupted by trapping at the surface.

## I. INTRODUCTION

In the description of both individual and collective dynamics of motile microorganisms a considerable simplification can often be achieved by partitioning their effect on the surrounding fluid into separate contributions from the organism body and the appendages —cilia or flagella—that confer locomotion. These contributions can further be simplified into those of equal and opposite point forces acting on the fluid, as required by the force-free condition on a free swimmer. Thus, peritrichously flagellated bacteria, with a bundle of rotating flagella aft of the body, are termed “pushers”, while the breast-stroke beating of paired algal flagella forward of the body defines a “puller” [1, 2]. Direct measurements of the flow fields around freely-swimming algae [3] and bacteria [4] have confirmed that the far-field flows are consistent with the singularity representation of swimmers.

The force dipole picture gives considerable insight into many features of swimming. Viewing microswimmer suspensions as a collection of interacting stresslets leads to an understanding [1, 5] of why a bacterial suspension exhibits “bacterial turbulence” [6–8] while a suspension of algae does not. At the individual level, the singularity picture explains accumulation of sperm cells at no-slip surfaces due to the reorientation of pushers to become parallel to such boundaries [9]. Attractive interactions between Stokeslets near a no-slip surface [10, 11] underlie the formation of “hydrodynamic bound states” of *Volvox* colonies [12], in which negatively buoyant chiral microswimmers are attracted together in the plane of the surface and orbit each other, a phenomenon later seen in several other systems [13–16].

Not surprisingly, the force dipole representation alone may fail to capture near-field effects, which may require higher-order singularities or the invocation of lubrication forces which become important in the near-field [17], an effect documented in swimming spermatozoa [18] and *E. coli* [19]. A clear breakdown of the stresslet picture is provided by interactions of the unicellular green alga *Chlamydomonas reinhardtii* with surfaces [20]. Whereas the stresslet approximation predicts that pullers nosedive into no-slip surfaces, experiments show instead an “inelastic scattering” phenomenon, where almost all incoming angles of swimming trajectories lead to approximately zero outgoing angle, corresponding to swimming parallel to the surface. In these experiments, the reorientation at the surface was shown to arise from direct ciliary contact interactions. Important later work [21] on scattering of *Chlamydomonas* by curved no-slip surfaces showed that similar geometry of reorientation can arise from hydrodynamic interactions without the need for direct contact with a wall.

The hydrodynamic interactions responsible for the rotation of algae away from the perpendicular orientation favored by the puller stresslet arise from the the undulatory beating of the two algal cilia. We term these “outboard” cilia, as they are displaced laterally on either side of the cell centerline. The time-averaged flow field around *Chlamydomonas* [3], shown in Fig. 1(a), as well as the time-resolved flow field [22], illustrates the swirling action of the flagellar beat, with an extended power stroke driving flows backward, and a contracted recovery stroke nearer the cell body, driving weaker flows forward. These complex time-averaged fields can be represented by a superposition of three Stokeslets: one for the cell body pointing forward, and one in the middle of each flagellum, pointing rearward, as in Fig. 1(b). The fully time-dependent problem can be modelled by time-varying combinations of singularities [24–26].

**FIG. 1.**
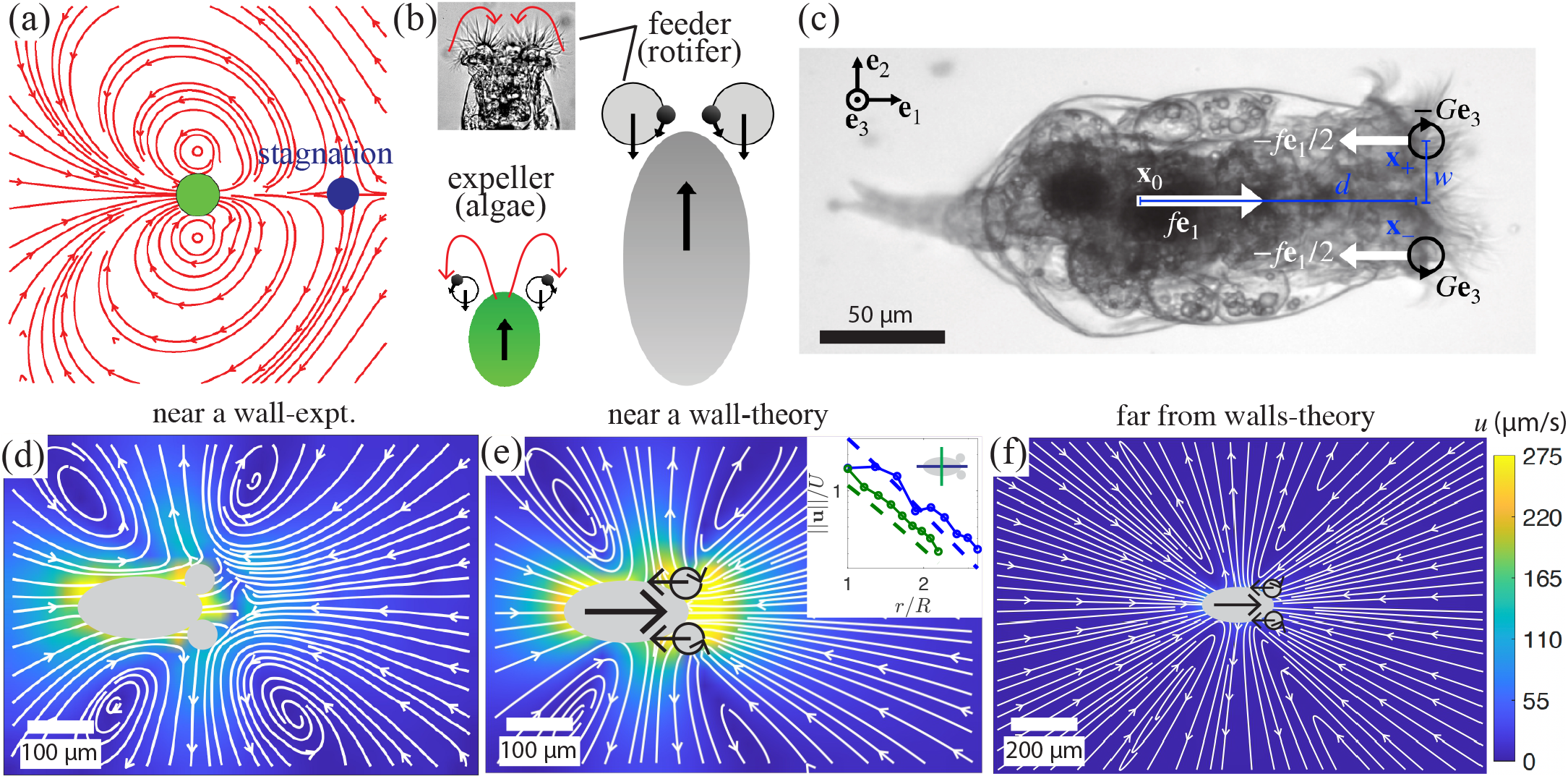
Flow fields of expellers and feeders. (a) Illustrative theoretical flow field for *Chlamydomonas reinhardtii* using a superposition of three Stokeslets and two rotlets (not fitted). (b) Schematics showing outward ciliary movement of the expeller and inward ciliary movement of the feeder. (c) Schematic representation of the body material frame and singularities employed in (2). The body frame consists of **e**_1_ (aligned with the body axis), **e**_2_ (in the plane of the coronae, pointing in either direction) and **e**_3_ = **e**_1_ ×**e**_2_ (pointing out of the page). The effect of the body on the fluid is represented by means of a system of Stokeslets and rotlets. The body Stokeslet of strength *f* **e**_1_, representing the drag on the fluid, is balanced by two −*f* **e**_1_*/*2 Stokeslets at **x**_±_ = **x**_0_ + *d***e**_1_ ±*w***e**_2_. The circulating flow induced by the cilia is modelled as two rotlets of strength ∓*G***e**_3_ placed at **x**_±_. (d) Magnitude and streamlines of the rotifer flow field. (e) Fitted approximation to (d), along with log-log plot of the velocity decay along (blue) and perpendicular to (green) the body axis; arrows schematically denote the orientation of the Stokeslets and rotlets. (f) Magnitude and streamlines of theoretical flow field for a rotifer a distance from the wall 10^4^ times larger than its fitted value in (e); the topology of the flow changes with only the trailing stagnation points surviving [23].

The notion that the flow field arising from the beating strokes of eukaryotic cilia can be represented by a Stokeslet appears as well in studies of so-called “mosaic” ciliated tissues [27], such as the epidermis of developing amphibians, in which a sparse population of multiciliated cells exists in a background of nonciliated cells. In the simplest picture, the action of a large number of cilia, with no phase synchrony, in driving flow along the tissue surface can be quantitatively captured by a single point force parallel to the surface. The representation of the flow due to a flagellum as that of a moving Stokeslet also forms the basis for a very large amount of work on the synchronization of cilia [28]. Yet, it is also intuitively reasonable that the orbits of the flagella, with the extended power stroke and contracted recovery stroke, could also be modelled as a point torque on the fluid, or a rotlet. Importantly, the flow fields for parallel orientation of both Stokeslets and rotlets near a no-slip surface both decay as 1*/r*^2^ [10, 29], so there is no way from the far-field decay alone to prefer one representation over another. Recent work has, however, shown that the time-averaged flow generated by a beating cilium near a wall is better represented as a rotlet, especially for near-field transport [30].

Here we reconsider the problem of microswimmers with outboard cilia in light of the background summarized above. Our primary observation is that Nature presents us with two broad classes of such organisms, distinguished by the direction of swirling flows created by the cilia. As in Fig. 1(a), the breaststroke beating of biflagellates such as *Chlamydomonas* sweeps fluid *away* from the cell apex; we name these “expellers”. By contrast, the corona of cilia in more complex multicellular organisms such as the rotifer *Brachionus plicatilis* shown in Figs. 1(b,c) directs flow *toward* the mouth, and are naturally termed as “feeders”. Rotifers are complex “animalcules” with internal organs and a nervous system, and serve as model organisms for a wide range of biological processes, from evolution [31] to aging [32]. The first observation of rotifers is variously attributed [33] to Antony van Leeuwenhoek [34] and John Harris [35], with decisive descriptions due to the former in a series of papers in the early years of the 18^th^ century [36–38]. Even in these very early works there is reference made to the tendency of rotifers to attach strongly to surfaces, which in light of the observations of surface scattering of *Chlamydomonas* serves to illustrate the fundamental distinction in surface interactions between expellers and feeders.

After outlining in Sec. II the experimental methods used here to study the swimming and surface interactions of rotifers, we present in Sec. III a quantitative analysis of the flow field around a freely-swimming rotifer and its representation in terms of a superposition of Stokeslets and rotlets. This leads naturally to consideration of the interactions of rotifers with no-slip surfaces, considered in Sec. IV, where we show that they exhibit a rapid transition from freely swimming to surface attachment. This can be understood quantitatively through a model akin to the stresslet one in which the additional contribution from the outboard cilia is represented by a rotlet doublet. The linear stability problem of such “composite” swimmers near a surface is studied in Sec. V. Finally, Sec. VI examines trajectories on larger spatial and temporal scales. We show experimentally that rotifers swimming near a surface exhibit the phenomenon of “run-and-stick”, in which roughly straight line swimming is interrupted stochastically by trapping at the surface through the mechanism discussed in Sec. V. A simple model of stochastic switching between bound and free states, similar in spirit to one used to study analogous transitions in *E. coli* [39], is shown to capture the essential features of the observations. The concluding Sec. VII highlights possible future directions of this research. Various details of calculations and data analysis are collected in Appendices A-D

## II. EXPERIMENTAL METHODS

We use the rotifer *Brachionus plicatilis* (strain 5010/4) as a model feeder organism [40]. It is approximately 210 *µ*m in length and 90 *µ*m in width, with individual cilia of length ∼50 *µ*m and beat frequency ∼20 −30 Hz. The cells can attain swimming speeds of 200 −400 *µ*m/s. They were grown in a coculture with the alga *Dunaliella tertiolecta* (strain 19/7c) as a food source in marine f/2 medium at 20 ^°^C, under a diurnal cycle of 12 h cool white light (∼15 *µ*mol photons/m^2^s PAR) and 12 h in the dark. All strains and media concentrates were sourced from the Culture Collection of Algae and Protozoa (CCAP) [41]. To isolate *B. plicatilis*, the coculture was passed through a 70 *µ*m diameter membrane filter (pluriStrainer) to remove the algae.

*B. plicatilis* is an invertebrate, with a dense ciliary array at the cell apex and a tail at the posterior end as shown in Fig. 1(c). While swimming, it retracts its tail, appearing like a prolate ellipsoid, and the cilia in front organize into two clusters of biaxial symmetric metachronal bands on either side of the cell axis, sweeping fluid towards the centrally located mouth. To quantify flow fields, we acquired brightfield images of *B. plicatilis* in f/2 medium infused with polystyrene tracer particles (mass fraction 0.01 %). Images were captured at 500 fps using a ×10 objective with a high-speed camera (Phantom V311) mounted on a Nikon TE-2000U inverted microscope. A dilute suspension of rotifers (100 cells cm^−3^) mixed with tracer particles was transferred into a chamber formed by two coverslips separated by a double-sided tape of thickness 4 mm. Fluid velocimetry using PIVlab was performed to analyze the flows [42]. The rigid boundary helps the rotifer swim parallel near to the surface long enough to capture high-resolution images of its flow-field. When far from the surface, image capture is difficult because the swimmers frequently move out of the focal plane, causing their bodies to appear defocused.

## III. FLOW FIELDS

Figure 1(d) shows time-averaged experimental flow-field of *B. plicatilis* swimming at a speed of 380 *µ*m s^−1^ nearly 350 *µ*m away from the upper surface of the chamber (See also SM Video 1 [43]). Assuming a typical rotifer size of *L* = 210 *µ*m, a maximum swimming speed of *U* = 400 *µ*m/s, and a kinematic viscosity of *ν* = 10^6^ *µ*m^2^/s, the Reynolds number does not exceed Re = *UL/ν* ∼ 0.08. We therefore work in the inertia-free limit Re = 0. We endow the swimmer centre of mass **x**_0_ with an orthonormal body-fixed frame {**e**_1_, **e**_2_, **e**_3_}, where **e**_1_ is the swimming direction and **e**_2_ is the normalised displacement between the cilia bundles located at

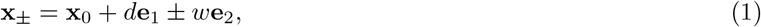

as in Fig. 1(c). The experimental flow in Fig. 1(d) was fit to a superposition of three Stokeslets of strengths *f* **e**_1_ (body drag), −*f* **e**_1_*/*2, −*f* **e**_1_*/*2 (thrust) located at **x**_0_, **x**_±_, as well as two rotlets with strengths ∓*G***e**_3_ at **x**_±_, representing the sweeping ciliary flow towards the mouth [17, 26] (Fig. 1(c) and Appendix A),

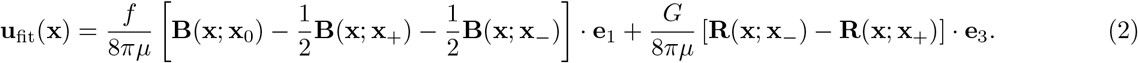

Here **B** and **R** are respectively the Blake tensors for a point force and torque near a no-slip wall [10]. The fit yields *f* ∼400 pN, *G*∼ 4 pN mm, *U* ∼400 *µ*m s^−1^, *d* ∼ 130 *µ*m, *w*∼40 *µ*m. The rotlets displacements *d* and *w* are comparable to the semi-axes, reflecting the bundles’ locations. Furthermore, for a bundle size of 25 − 50 cilia, the force per cilium is on the order of 4 −10 pN, consistently with the literature [44]. A positive value of *G* makes the organism a “feeder”, as in Fig. 1(b). The torque exerted by a single bundle should be on the order of (thrust per cilium) × (bundle circumference). Assuming a thrust of 10 pN per cilium and a bundle radius of 25 *µ*m, the estimate *G* ∼2 pN mm resembles the fitted value. Finally, neglecting the no-slip wall and treating the body as a prolate ellipsoid, the swimming speed in an unbounded fluid is related to the thrust by *f* = 6*πµbζU*, where [45]

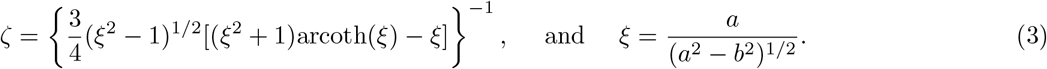

Taking the semi-axes to be *a* ∼100 *µ*m, *b* ∼ 50 *µ*m and assuming *µ* = 10^−3^ Pa s, a thrust of 400 pN should result in a swimming speed *U* ∼ 350 *µ*m s^−1^, similar to the experimental value. The fitted flow in Fig. 1(e) displays good quantitative agreement with the experimental data in Fig. 1(d) up to 1.5 body lengths ahead of the swimmer. The flow topology of rotifers changes far from the chamber wall, where the two elliptic stagnation points forward of the bundles Fig. 1(d,e) disappear leaving only the trailing stagnation points [4, 23, 46]. Notably, unlike for *C. reinhardtii* in Fig. 1(a) [3], the rotifer flow field in the absence of a boundary (Fig. 1(f)) lacks a stagnation point ahead of the body, owing to the smaller aspect ratio and the opposite circulation near the cilia bundles. The inset in Fig. 1(e) suggests a decay rate *r*^−1.43^ for the experimental flow perpendicular to the body axis and a *r*^−1.78^ decay along it. These are consistent with *r*^−2^ stresslet flow, which is unaffected by the presence of the wall within a distance *r* ≪ 2*h*∼ 720 *µ*m of the singularities, with *h* being the fitted height above the chamber wall. For *r* ≫ 2*h*, the presence of the wall is felt and the flow decays like *r*^−3^, rather than *r*^−2^. Interestingly, very strong confinement by two no-slip surfaces has been observed to reverse the sense of circulation of the vortices straddling the body of *Chlamydomonas*, turning an expeller flow into one that resembles a feeder [47].

## IV. SCATTERING FROM A NO-SLIP SURFACE

While the interactions between pusher swimmers and boundaries have been shown to be chiefly hydrodynamic [9], as discussed in the introduction, there is evidence that pullers can interact with walls both through direct wall contact [20, 21] and through the fluid [21, 26]. Moreover, while the leading-order stresslet flow suggests that pullers should nosedive into the wall, both scattering [20, 21] and trapping are observed experimentally. Here we study the hydrodynamic picture of the different scattering properties of feeders and expellers, exemplified by *B. plicatilis* and *C. reinhardtii*. We employ a far-field expansion that includes both the force dipole term [48], and the rotlet dipole that arises from contracting the two opposite rotlets into a single singularity. This approach offers a simple framework that can be incorporated into simulations of many-swimmer systems and coarse-grained continuum theories [1].

We quantify the strength of the rotlets via the signed “feeder number”

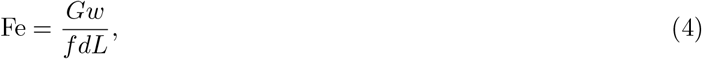

where *L* is a again the typical swimmer size. We shall refer to swimmers with |Fe| ≫ (≪)1 as “strong (weak) feeders/expellers”. *B. plicatilis* is a weak feeder (Fe∼0.06) while *C. reinhardtii* is a strong expellers with Fe ∼−5 (using *L* ∼5 *µ*m, *d* ∼5 *µ*m, *w* ∼10 *µ*m, *f*∼ 7.2 pN, *G*∼− 87 pN *µ*m, *U*∼ 100 *µ*m s^−1^, and *e* ∼ 0.75 [3, 44, 49]). The rotlet flow becomes comparable to that of the force dipole at the “feeder length”

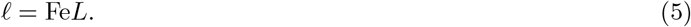

Indeed, |𝓁| ∼5*L* matches the *C. reinhardtii* stagnation point [3], located ∼ 6 radii ahead of the cell body. In the far field *r* := ∥**x**∥ ≫ *L*, a swimmer may be described as a sum of flow singularities, specifically a force dipole of magnitude 𝒪(*fd/µr*^2^), and higher-order gradients of Stokeslets of strength 𝒪 (*fd*^2^*/µr*^3^) or 𝒪 (*fw*^2^*/µr*^3^) [3, 17], a rotlet dipole of strength 𝒪 (*Gw/µr*^3^) and asymptotically smaller terms that we neglect. Since *d, w*∼*L*, the force dipole dominates the quadrupole for *r* ≫*L*. We note here that if *G*∼ *f L* as we may expect, then the rotlet dipole 𝒪 (*Gw/µr*^3^) 𝒪 ∼ (*f Lw/µr*^3^) is dominated by the force dipole for *r* ≳ *L*. In practice, however, the relation *G* ∼*f L* only applies to the peak force and torque *f* ^*^, *G*^*^. Because only a fraction of the peak force is converted into thrust (0.25 for *C. reinhardtii*), the average force *f* which sets the swimming speed [44] is typically much lower (*f* ∼ 0.25*f* ^*^ for *C. reinhardtii*). Therefore, while *G*^*^*w/f* ^*^*dL* ∼ −2, the “effective” feeder number is Fe ∼ −5 for *C. reinhardtii*. For strong feeders and expellers the dipole is therefore comparable to the rotlet dipoles for 1 ≪ *r/L* ≪ |Fe|. This validates representing the swimmer as a force dipole and torque dipole term of respective strengths *fd* and *Gw* centred at the same point **x**_0_. Despite the singularity approximation breaking down near a wall, it has been shown to qualitatively reproduce near-field dynamics [4, 9, 17, 50].

The wall modifies the trajectory of the swimmer through an image flow **u**^*^ ensuring no-slip. The translation of a prolate ellipsoid is governed by the Faxén equation,

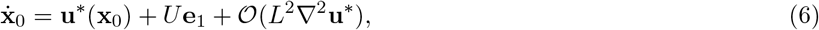

while the orientational dynamics is expressed by Jeffery’s equation [17],

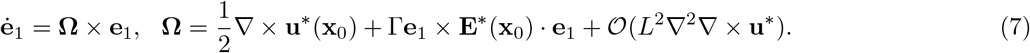

Here, **E**^*^ = (∇**u**^*^ + ∇^T^**u**^*^)*/*2 is the image rate-of-strain tensor, and Γ= (1 − *e*^2^)*/*(1 + *e*^2^) is the shear-alignment parameter with *e* = *b/a* the aspect ratio. We henceforth neglect the asymptotically sub-leading terms in (6) and (7). We now apply the above to the scattering of microswimmers from a no-slip surface, focusing first on expeller dynamics exemplified by *Chlamydomonas*. For context, Fig. 2(a) shows a typical scattering event of *C. reinhardtii* in which one cilium makes brief contact with the surface. Figure 2(b) examines the case in which purely hydrodynamic interactions govern scattering, obtained by numerical integration of the dynamics (6) and (7), using typical values *d* ∼5 *µ*m, *w* ∼10 *µ*m, *f* ∼7.2 pN, *G*∼ −87 pN *µ*m, *U* ∼100 *µ*m s^−1^, *e*∼0.75 [3, 44, 49]. For a range of incoming angles *θ*_in_ we observe gliding along the wall for a short distance followed by turning away from the wall with a scattering angle *θ*_out_ that is monotonically increasing with the incident angle. Figure 2(c) plots the scattering data of Contino, et al. [21] for the wild type strain CC125 and the short flagella mutant SHF1 of *C. reinhardtii* in the range *θ*_in_ ≳ 44^°^ where boundary interactions are mostly hydrodynamic. In order to compare the results of the singularity model with these data we computed the ⟨*θ*_out_⟩ for 10^3^ random values of the roll angle. The average value and spread associated with the roll angle are shown respectively as a solid line and shaded region in Fig. 2(c) indicate that this model accurately reproduces the approximately linear growth of ⟨*θ*_out_⟩ with *θ*_in_. Indeed, we fit ⟨*θ*_out_⟩ ∼0.69 · *θ*_in_ + 11.13^°^, while ⟨*θ*_out_⟩ ∼0.59 · *θ*_in_ + 22^°^ for the CC125 mutant and ⟨*θ*_out_⟩ ∼0.64 · *θ*_in_ + 21^°^ for the SHF1 mutant [21]. These results provide a quantitative validation of the model of *Chlamydomonas* as a swimmer governed by the sum of a puller stresslet and an expeller rotlet doublet.

**FIG. 2.**
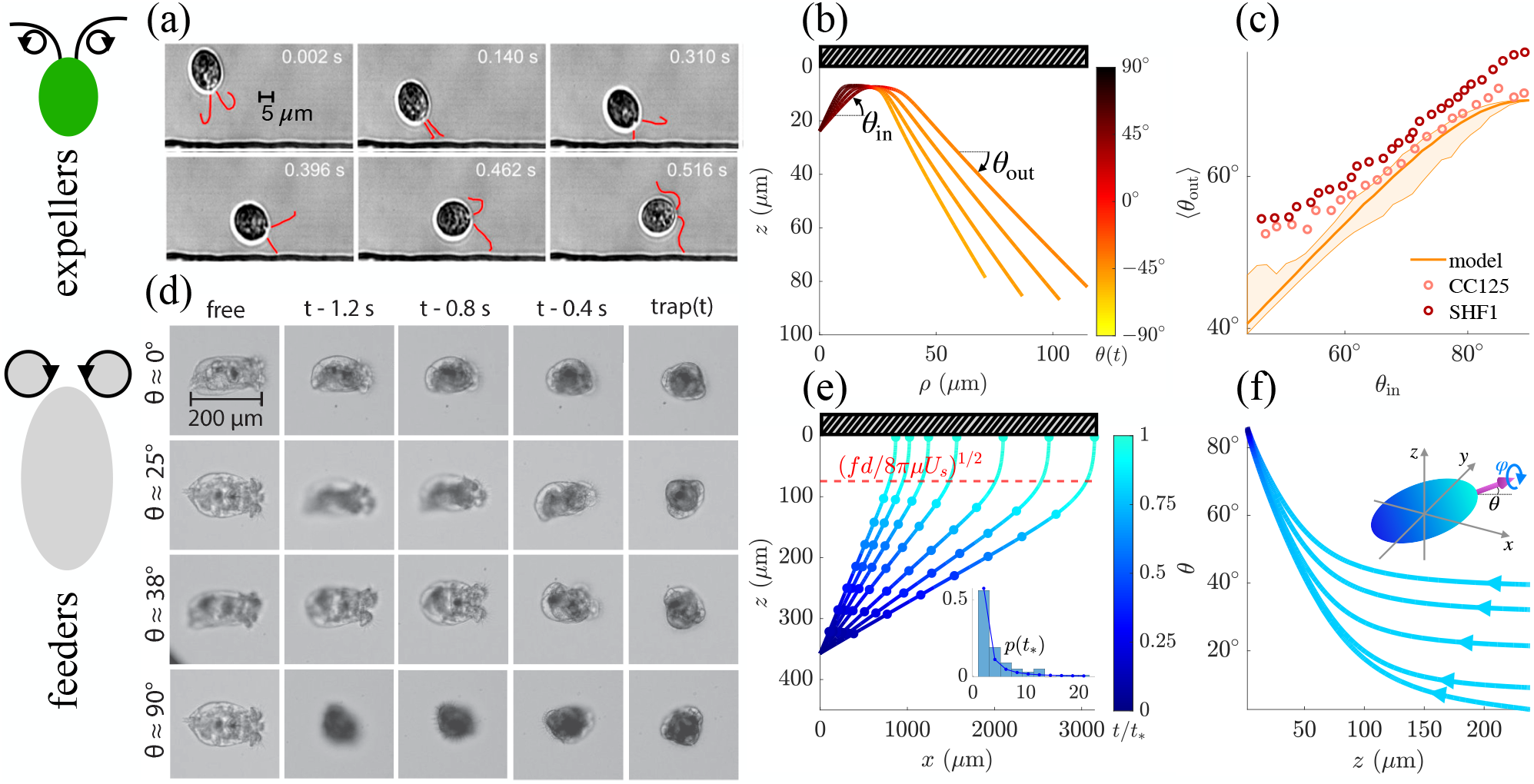
Scattering of expellers and feeders from surfaces. (a) Scattering of *C*.*reinhardtii* off a rigid boundary via steric interactions, from [20] with permission. (b) Computed trajectories appropriate to *C. reinhardtii*. (c) Average scattering angle ⟨*θ*_out_⟩ as a function of incident angle *θ*_in_ for *C. reinhardtii*. Orange curve shows the predictions of singularity model, with shaded region indicating variations associated with roll angle. Circles are the scattering data for the mutants CC125 and SHF1 from [21]. (d) Trapping of four rotifers with different initial pitches (see also SM Video 2 [43]). (e) Theoretical trajectories of rotifers show “snapping” to the boundary for a range of initial pitches and zero initial roll. The colorbar represents the proportion of the total time elapsed, while circles mark the position at regular time intervals, showing a speed increase near the boundary. The red line marks the distance *z*^*^ in (8) at which the dipole flow becomes comparable to the swimming speed. The inset shows the distribution of the impact time *t*_*_ and a fitted ‘ballistic’ p.d.f. 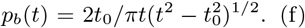. (f) Evolution of the pitch and roll *θ* and *ϕ* for five trajectories with different *θ*_0_ and *ϕ*_0_. Variations in the roll are correlated with the color shade. As the distance to the wall decreases, the pitch increases to 90^°^ and the roll changes abruptly near impact.

Turning to rotifers, Fig. 2(d) shows examples of their rapid “snapping” to a solid boundary. In the geometry of an inverted microscope, these are views from below as rotifers that are initially oriented with their long axis **e**_1_ parallel to the upper chamber coverslip turn and rapidly rotate to become attached to that surface, such that **e**_1_ points away from the observer. As remarked earlier, this phenomenon was known to van Leeuwenhoek, who said in his famous letter to the Royal Society of 17 October, 1687, *“*… *These Animals also had a second movement; for when they were unable to make any progress by swimming, they attached themselves to the glass by the organs at the front of the head; and then they drew their body up short*…*”* [51]. Our studies show that the process of snapping typically takes on the order of 1− 2 s from initiation to vertical alignment. Once attached, rotifers are observed to spin around **e**_1_ with a period of ∼5 s, sometimes for many complete rotations before ultimately detaching. This motion is likely related to the spinning motion around **e**_1_ seen during free swimming. We discuss the long-time statistics of the switching between swimming and sticking in Sec. VI below.

Using the fitted values of *fd* and *Gw* in the stresslet+rotlet doublet model, Fig. 2(e) shows that numerically obtained snapping trajectories recapitulate the behavior observed in Fig. 2(d). As the wall is approached the flow due to the strong puller stresslet increases as 1*/z*^2^, and thus we can define the lengthscale

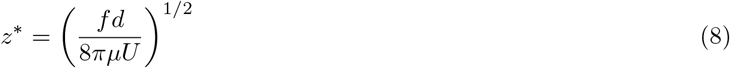

at which the stresslet flow is comparable to the swimming speed. From the angular dynamics in Fig. 2(f) we deduce that the body turning is driven by the strong puller stresslet. Defining the yaw *χ*, pitch *θ*, and roll *φ* of the body frame by {**e**_1_, **e**_2_, **e**_3_} = *R*_*z*_(*χ*)*R*_*y*_(*θ*)*R*_*x*_(*φ*), Fig. 2(f) shows that the pitch rapidly increases to nearly 90^°^ and impact is associated with abrupt twisting motion along the body axis.

## V. STABILITY ANALYSIS OF TRAJECTORIES

In this section we show that the presence of “feeder” rotlet flow turns a swimmer located at a distance *h* from a wall towards the no-slip wall when the trajectory is nearly perpendicular or parallel to the wall. Conversely, an “expeller flow” generally rotates the rotifer away from the wall. For a trajectory that is nearly perpendicular to the wall, i.e. with 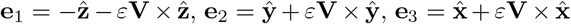, up to 𝒪 (*ε*^2^), a careful analysis given in Appendix C shows that the swimmer moves towards the wall with speed

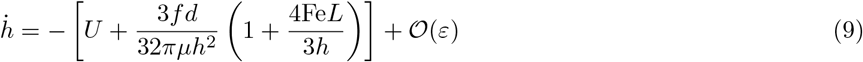

Therefore, feeders speed up as they approach the wall, while expellers slow down or even hover if 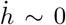. Rotlets control the snapping if the crossover length *h* ∼ Fe*L* = 𝓁 is much larger than the body size, i.e. if the organism is a strong feeder or expeller. Moreover, a nosediving puller is instantaneously rotated towards the wall if

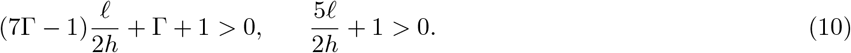

For large force dipoles (|𝓁*/h*| ≪ 1) the swimmer always aligns perpendicular to the wall [17], while for strong rotlet dipoles or near collision (|𝓁*/h*| ≫ 1) the swimmer tends to align parallel to the wall when *G <* 0 (expeller) and perpendicular to the wall when *G >* 0 (strong feeder) provided the body is not too spherical. This analysis reveals a fundamental difference between rotifers and green algae: whereas the former behave essentially like stresslets when colliding with the wall (and are thus stable), expeller algae feel the destabilizing effects of the rotlet and are thus unstable [see Fig. 2(b,c)].

Considering now initially parallel trajectories with 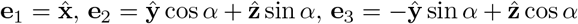, since the force dipole does not rotate the swimmer [17], the leading-order effect comes from the rotlets. The presence of the rotlet tends to steer the swimmer towards the wall for feeders and away from the wall for expellers. Explicitly, we find that the swimmer is rotated towards the wall if

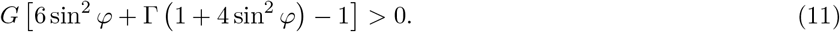

This is always the case for feeders provided *α* is not too close to a multiple of *π*. If this happens, vorticity prevails over shear alignment and the swimmer turns away from the wall. For expellers (11) predicts that the swimmer will tend to turn away from the wall provided sin *α* is not too small, which matches numerical findings [26].

The dynamics of trajectories can be quite complex as a consequence of the competition between the stresslet and rotlet dipole contributions. Figure 3 shows four trajectories for feeders and expellers initially oriented parallel to the surface. Fig. 3(c) shows two trajectories corresponding to parameters appropriate to *C. reinhardtii* with initial roll *φ*_0_ = 11.5^°^ and *φ*_0_ = 28.6^°^. For the former, just inside the stability region, the pitch initially increases and the cell moves towards the wall, until *φ* becomes too large. When this occurs, the organism rapidly rotates away from the wall (decrease in *θ*) and then departs, never to return. For *φ*_0_ = 28.6^°^, the alga rotates away from the wall and departs. Figure 3(d) instead shows two trajectories appropriate to *B. plicatilis* with *φ*_0_ = 5.7^°^ and *φ*_0_ = 51.6^°^. For the former, inside the unstable region, the rotifer swims away from the wall. When *φ*_0_ = 51.6^°^, the organism rotates towards the wall and crashes after a temporary increase in *z* due to the force dipole.

**FIG. 3.**
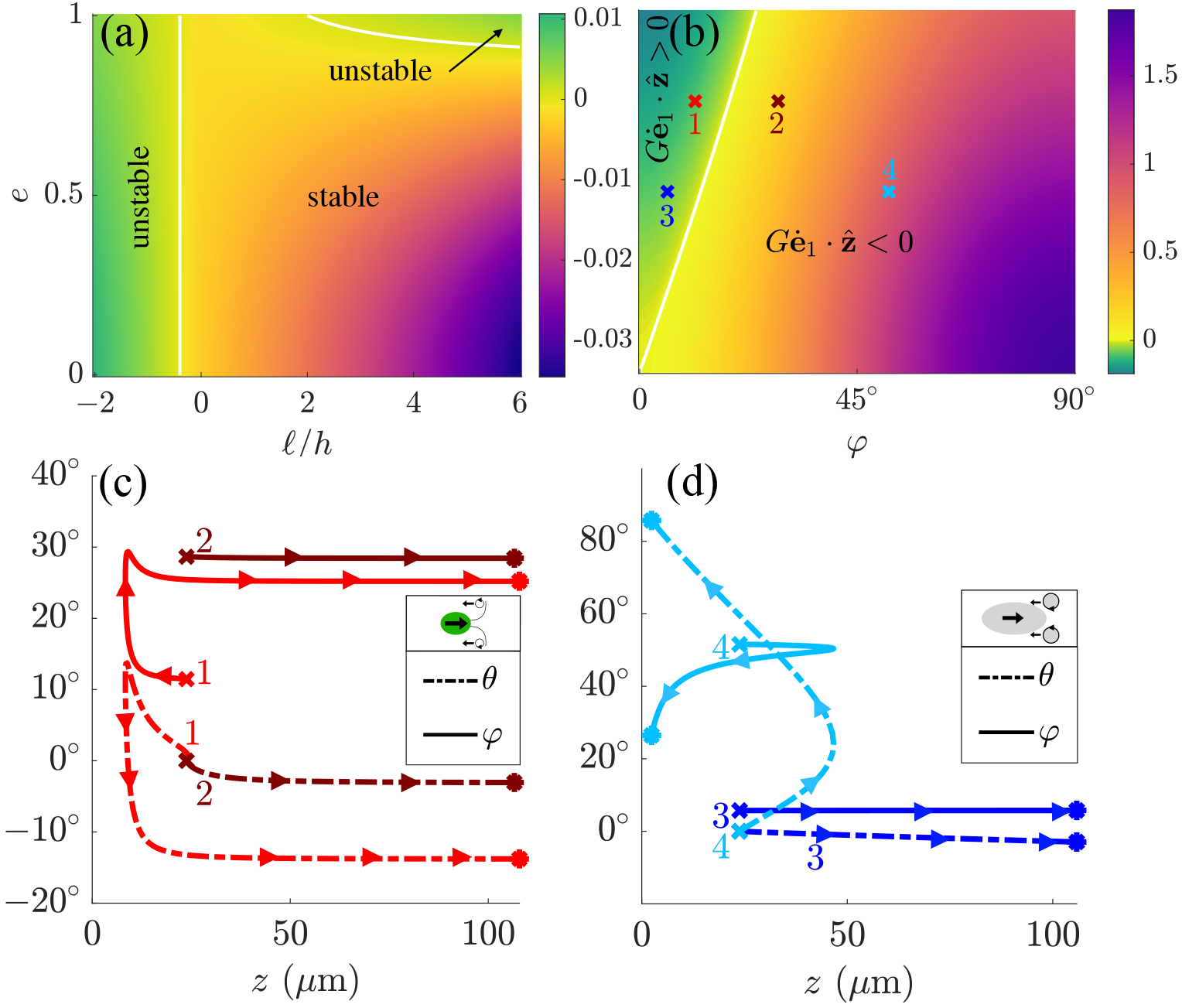
Results of stability analysis. (a) The case of a trajectory initially oriented nearly perpendicular to the surface. Trajectories were obtained with the value of *Fd* extracted from PIV, *U* = 0, *h* = 50 with a nearly vertical swimming direction, *t*_max_ = 40 and a 100 × 100 grid. The aspect ratio *e* and dimensionless feeder length 𝓁*/*2*h* were varied, and the plot shows max_*i*_[|*e*_*i*_(*t*_max_)|− |*e*_*i*_(0)|] for *i* ∈ {1, 2}. White lines mark the theoretical boundaries of the stability regions. (b) Stability diagram of a rotlet dipole with 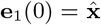 sand roll *φ*. For *G >* 0 (feeder) the dipole is rotated towards the surface 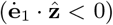 for *φ* sufficiently large, while when *G <* 0 (expeller) the dipole rotates away from the surface unless *φ* is small. (c) Sample trajectories for representative parameter values of *C. reinhardtii* in different parts of phase space. (d) Sample trajectories for representative parameter values of rotifer in different parts of phase space.

## VI. MOTILITY STATISTICS NEAR SURFACES

The transitions back and forth that rotifers exhibit between free swimming parallel to a surface and to attaching to it lead to an intriguing type of random walk loosely analogous to several other multi-mode locomotion motifs: the “run- and-tumble” locomotion of bacteria such as *E. coli* [52], the “run-reverse-flick” locomotion of *Vibrio alginolyticus* [53], and the “run-and-turn” locomotion of *Chlamydomonas* [54], and the “run-stop-shock” locomotion of *Pyramimonas octopus* [55]. It is perhaps most closely related to the behavior exhibited by a particular pathogenic strain of *E. coli* that stochastically transitions between runs parallel to surfaces and adhesive events [39]. We analyzed 62 tracks of *B. plicatilis* with typical durations ranging of 20 s. Figure 4(a) shows typical examples of those trajectories, which follow what we term a “run-and-stick” sequence. As detailed in Sec. IV, a rotifer that is close to a surface may rapidly reorient to become perpendicular to the surface, becoming stably attached, as in Fig. 2(d). While the rotifer’s strong puller stresslet flow field leads to strong attachment, they do eventually release owing to biological activity (SM Video 3 [43]). While attached, the aforementioned spinning of rotifers around the body axis **e**_1_ leads to a randomized swimming direction upon their release (SM Video 1 [43]).

**FIG. 4.**
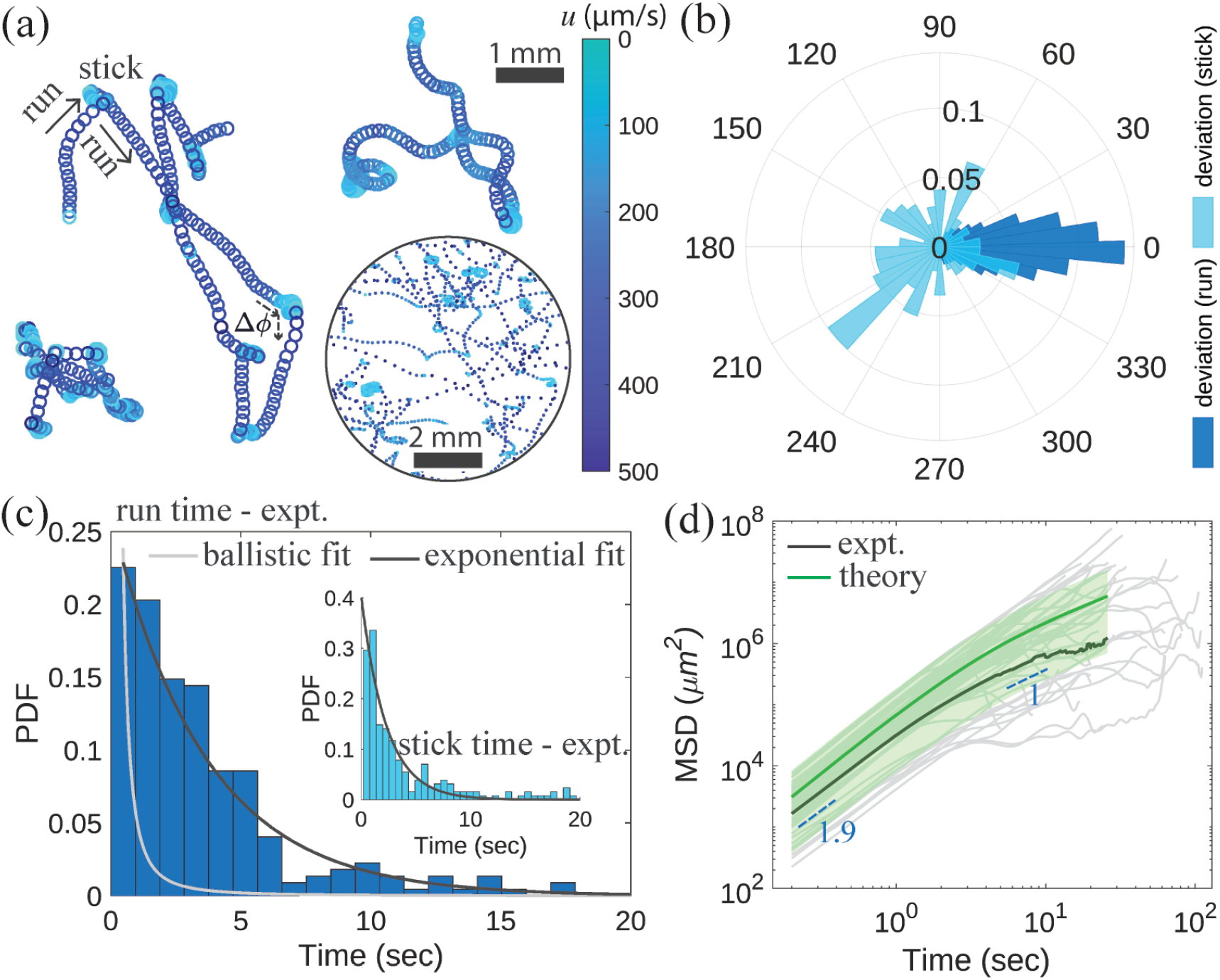
Motility statistics of swimming rotifers. (a) Examples of trajectories near the top chamber surface, color-coded by the swimming speed. (b) PDF of ‘run deviations’ between consecutive stick events and ‘stick deviations’, the difference between incoming and outgoing angles. (c) PDF of the run-time and stick-time durations. (d) MSD (gray - individual tracks, black - average) of *B. plicatilis* exhibits a transition from super-diffusive to diffusive behavior, owing to sticking near the surface. The green curve and shading illustrate the predicted MSD and 2 standard deviations based on theoretical calculations.

We quantify the run-and-stick trajectories by examining first in Fig. 4(b) two measures of angular deviations: (i) the ‘run-deviations’ Δ*θ*_*i*_ = *θ*_*i*_ −⟨*θ*_*i*_⟩ of trajectory directions *θ*_*i*_ from the mean direction ⟨*θ*_*i*_⟩ of the *i*^th^ swimming event, sampled in steps of 0.2 s, and (ii) the ‘stick-deviations’, the angular change Δ*ϕ* between incoming and outgoing directions at each sticking event. A sharp peak around 0^°^ for run-deviations suggests that the cells swim mostly in a straight line until reoriented by stick events. The role of biological activity in sticking and unsticking events becomes even more evident once we assess the ‘run time’ and ‘stick time’ histograms in Fig. 4(c). Averaged over all 62 analyzed tracks near the top surface we find ⟨run time⟩ = 3.6 s and ⟨stick time⟩ = 5.5 s, with both distributions decaying exponentially, signifying a Poisson processes. Such a process is consistent with the picture that the organism swims roughly parallel to the wall until its pitch stochastically fluctuates sufficiently to trigger a snapping event. In this case the waiting time between events should be exponentially distributed with probability density function (p.d.f.) *p*_*e*_(*t*) = *λe*^−*λt*^. An alternate view on the waiting time distribution is that the organism swims ballistically at all times except close to snapping and that all the randomness in the trajectory is embedded in the initial pitch *θ*. In this case we can evaluate the corresponding ‘ballistic’ impact time *t*_*_ for an incidence angle *θ* as *t*_*_ ∼ *t*_0_*/* sin *θ*, where *t*_0_ = *h/U* is a typical swimming time. Because the transition to sticking occurs rapidly relative to the swimming timescale, it can be ignored. If the incidence angle is chosen uniformly at random in [0, *π/*2], the ‘run time’ distribution follows the ballistic p.d.f. 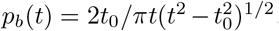. Going back to Fig. 2(e), we test this approximation by fitting *p*_*b*_ to the p.d.f. of the collision time for 100 trajectories with uniformly distributed initial pitches and an initial height *h* = 360 *µ*m (inset). Results show excellent agreement with *χ*^2^ error ∼ 0.02.

In order to see which of the exponential or ballistic p.d.f. better accounts for the data, we fitted both *p*_*e*_ and *p*_*b*_ to the data (without automated binning) and computed the *χ*^2^ error in the fit. For *p*_*e*_, this was ∼0.15, while for *p*_*b*_ it was ∼0.74. This analysis shows that snapping is explained much more convincingly by swimming noise rather than ballistic motion with initially random angles. From the exponential fit, we find a mean free-flight time of ∼ 3.6 s.

Next we examine the mean squared displacement (MSD) of rotifers undergoing run-and-stick locomotion. Figure 4(d) shows the ensemble of MSD measurements for the analyzed tracks and the ensemble average MSD. There is a clear transition from near-ballistic motion for short times to a diffusive regime on longer timescales, with a crossover time of ∼2−4s. To explain this, we propose a mean-field model of a population of rotifers near a boundary in which the population is coarse-grained into a local number density of freely swimming and trapped rotifers.

Let *f* ^+^(**x**, *θ, t*) be the number density of freely swimming rotifers at position **x** and angle *θ* at time *t*. Similarly, let *f* ^−^(**x**, *t*) be the number density of trapped rotifers at position **x** and time *t*. Based on the motility statistics in Fig. 4, we assume that individual rotifers transition between freely swimming and trapped at a rate *ν*_−_ and from a trapped state to freely swimming at a rate *ν*_+_. Neglecting diffusion between trapping events on account of the small angular displacement in Fig. 4(b), we propose that the distributions obey the evolution equations

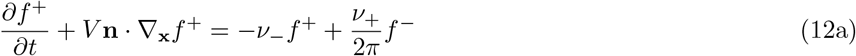

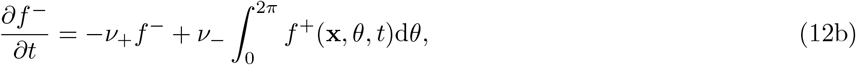

where **n** = [cos *θ*, sin *θ*]. We term (12a) and (12b) a *run-and-stick model*. Eq. 12a is the statement that a parcel of freely swimming rotifers with angle *θ* loses *ν*_−_*f* ^+^ swimmers per unit time due to sticking, and gains *ν*_+_*f* ^−^d*θ/*2*π* swimmers per unit time as a result of unsticking events. The 1*/*2*π* factor denotes the fact that the swimming angle upon release is uniformly random due to loss of orientation in the trapped state. Likewise, Eq. 12b models the fact that the trapped population loses members at a rate *ν*_+_ and gains members (with arbitrary swimming angle) at a rate *ν*_−_. From *f* ^+^ and *f* ^−^ we may define the total rotifer number density *ρ*(**x**, *t*) as

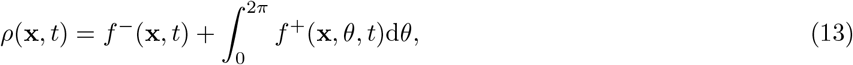

which, from (12a) and (12b), satisfies the conservation law

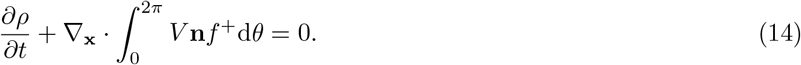

Appendix D provides the details of the calculation of the MSD of the run-and-stick model, with the result

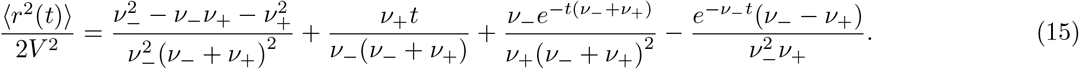

At short times 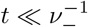, there is ballistic behavior with ⟨*r* (*t*)⟩ = *V*^2^*t*^2^ + 𝒪(*V*^2^*ν* _−_*t*^3^), crossing over to linear behavior_−_ at long times, from which we calculate an effective diffusion coefficient for the population based on the asymptotic result MSD ∼ 4*Dt* for *t* → ∞ in two dimensions (Appendix D), yielding

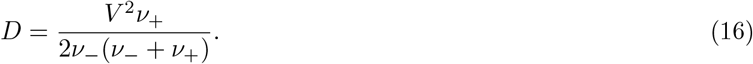

Taking *V* = 280 *µ*m s^−1^, *ν*_+_ = (5.5 s)^−1^, *ν*_−_ = (3.6 s)^−1^, we obtain *D* ∼ 0.057 mm^2^ s^−1^, in good agreement with experiments.

The result (16) can be compared with the classic run-and-tumble (RT) process, which formally corresponds to the limit *ν*_+_ → ∞. This yields the quasi-steady result ∂*f* ^−^*/*∂*t* = 0, and hence

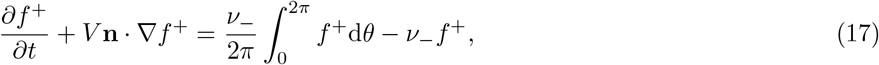

which corresponds to an RT process with frequency *ν*_−_ [56]. In this limit, *D* ↗ *D*_RT_ = *V* ^2^*/*2*ν*_−_; for finite *ν*_+_, *D < D*_RT_ so the population spreads more slowly than for RT due to the extra latency from sticking events. Equivalently, there an effective free-flight time smaller than the inverse sticking rate,

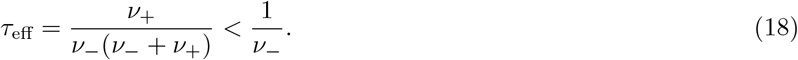

## VII. DISCUSSION

In this paper, we demonstrated that puller microswimmers may be classified as “feeders” or “expellers” depending on the sense of circulation of the cilia-driven flows. These two classes may be identified by their near-field flows (Fig. 1), with expellers exhibiting a stagnation point ahead of the cell body and feeders conversely presenting incoming flow at their apex. Boundary interactions also differ in relation to the “feeder number” Fe quantifying the relative strengths of the Stokeslet and rotlets associated with propulsion. While strong expellers (|Fe| ≫ 1) glide along the wall and then depart, feeders tend to collide and even attach to the boundary. By comparing with experimental data, we demonstrated that such behaviour is closely captured by an approximation consisting of a force dipole and a rotlet dipole located at the same point within the organism body (Fig. 2). A linear stability analysis confirms that the rotlet dipole generally turns feeders towards the boundary for both parallel and orthogonal incoming trajectories, leading to collision, while expellers are rotated away from the boundary, leading to scattering (Fig. 3). Motility statistics (Fig. 4) reveal that both the free-flight and the trapped times of rotifers are exponentially distributed, signifying that sticking and unsticking are well described by Poisson processes. While rotifers perform nearly ballistic motion on timescales much shorter than the average free-flight time, their long-term motion is significantly modified by sticking events. In particular, the changes of incoming and outgoing directions are random, leading to a crossover from ballistic to diffusive scaling. The motility near the surface is well-described by a mean field run-and-stick model predicting an effective diffusive behaviour in good quantitative agreement with experimental data.

While the results presented here show that many aspects of the swimming dynamics of rotifers can be understood using familiar methods in fluid mechanics, we note that rotifers possess muscles with which they can deform their body and sensory organs for touch and light. They are therefore capable of significantly more complex behaviors than the bacteria and protists that serve as paradigms of microswimmers. Thus, their dynamics near surfaces may reflect at least in part a tactic response as much as a purely passive hydrodynamic phenomenon. Finally, the possibility of interesting collective effects, whether in bulk or at surfaces, from organisms such as rotifers remains to be explored.

## ACKNOWLEDGMENTS

We thank Rebecca Poon and Francesco Boselli for assistance with PIV, Kyriacos Leptos for numerous discussions and Brian Ford for historical background on rotifers. This work was supported in part by Grant No. 7523 from the Gordon and Betty Moore Foundation, and the John Templeton Foundation.

## Appendix A: Fitting the flow

The rotifer’s body is seen to lie within the focal plane throughout the analyzed videos, implying that it is parallel to the no-slip wall at *z* = 0. We thus orient the PIV field of view so that the body frame of reference is 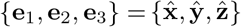 and fit the flow in the *xy* plane. Such procedure is performed after removing the solid-body motion velocity corresponding to the rotifer location. We model the rotifer via a far-field approach whereby the body and locomotion apparatus are replaced by point singularities; the thrust is taken to be being parallel to the body axis, and the balancing effects of the cilia bundle and the body drag are represented by Stokeslets of strength 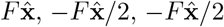 located at **x**_0_, 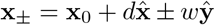. We additionally place two rotlets of strengths ∓*G***e**_3_ at **x**_±_, as in Fig. 2. Because the field of view is confined to the *xy* plane, we can only detect the 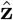 component of the rotlets from the cilia bundles. The flow Ansatz is then given by Eq. 2 in the main text, where **B** is the Green’s function for a point force near a no-slip boundary,

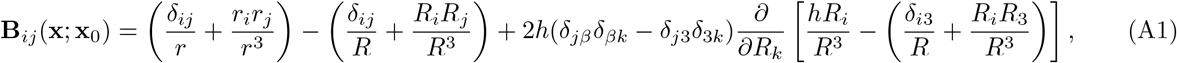

with *β* = 1, 2, 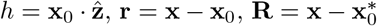 and 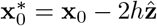. Similarly, **R** is the Green’s function for a point torque near a no-slip boundary,

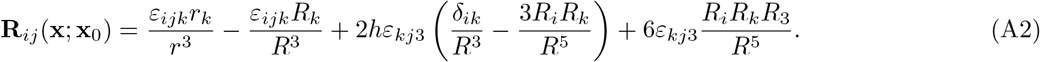

For simplicity, we pin **x**_0_ on the body axis, so we only fit the distance from the wall and the axial position of the body Stokeslets, giving 6 fitting parameters in total.

In fitting the flow field, we care particularly about capturing the four near-field lobes, a signature of proximity to a no-slip surface. We therefore propose as the metric to minimize the function

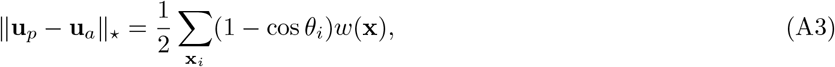

where **u**_*p*_ and **u**_*a*_ are the PIV and analytical flows, *θ*_*i*_ is the angle with respect to the PIV flow direction and the **x**_*i*_ are the PIV lattice nodes after removal of the region corresponding to the rotifer’s body. The weight function *w*(**x**) is the sum of four Gaussians centred at the vortices,

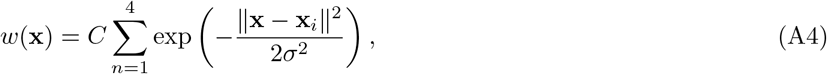

where *σ* is the standard deviation (set by the vortex size), assumed for simplicity to be equal for all vortices, and *C* is a (numerically determined) normalising constant such that

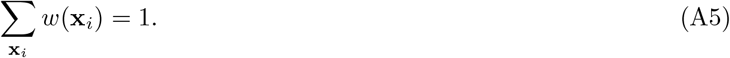

We chose *σ* = 1 in our fitting and constrained the singularity locations lie close to the solid-body region of PIV, specifically within a rectangle 1.5 times larger than the rotifer’s body. This fit only determines the flow up to rescaling *f* → *κf, G* → *κG*. We find the optimal value of *κ* via a least-squares method.

## Appendix B: Equations of Motion

In the absence of a wall, the swimmer swims without rotation in a straight line with velocity *U* **e**_1_. The no-slip condition on the wall induces a perturbation flow **u**^*^(**x**) that may formally be obtained by placing suitable “image singularities” at the mirror-image of the body’s location 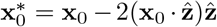 in order to exactly cancel the organism’s flow at the wall. Such a flow advects and rotates the organism according to the Faxén laws for a prolate ellipsoid,

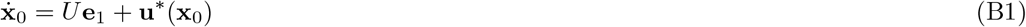

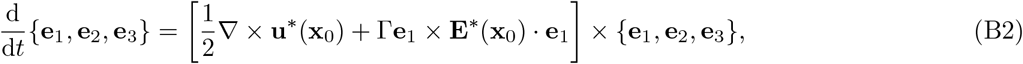

where Γ= (1 −*e*^2^)*/*(1 + *e*^2^) is the Bretherton parameter encoding shear-alignment, 0 ≤ *e* ≤ 1 is the ellipsoid’s aspect ratio, and **E**^*^ = (∇**u**^*^ + ∇^T^**u**^*^)*/*2 is the rat-of-strain tensor of the image flow. For simplicity, we henceforth model the rotifer as a superposition of a force dipole of strength *fd* along **e**_1_ and a rotlet dipole of strength *Gw* along **e**_2_.

The image flow 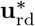 induced by the rotlet dipole (located at **x**_0_) at an arbitrary point **x** is

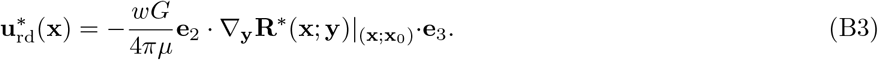

The image flow **R**^*^(**x**; **y**) · **e**_3_ generated by a rotlet of strength **e**_3_ located at **y** is

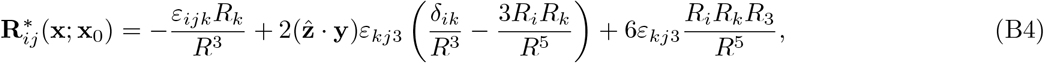

where **R** = **x** − **y**^*^, 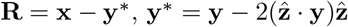. The vorticity and rate-of-strain tensor of this flow at the position **x**_0_ are

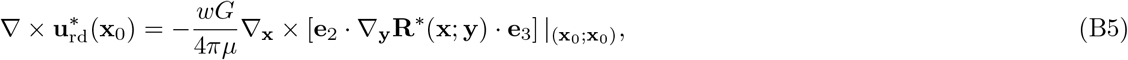

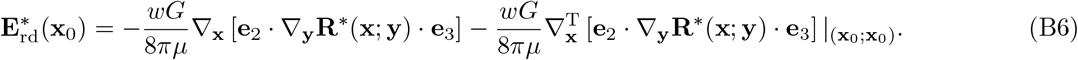

On the other hand, the image flow 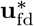 produced by the force dipole (located at x_0_) at an arbitrary point x is

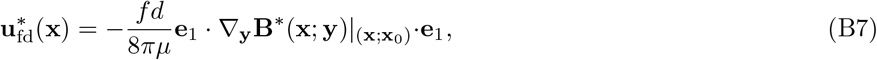

where the image flow **B**^*^(**x**; **y**) · **e**_1_ generated by a Stokeslet of strength **e**_1_ located at **y** is

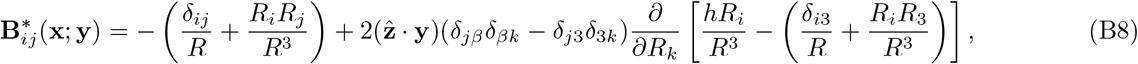

where **R** is defined as in (B4). The corresponding vorticity and rate-of-strain tensor are

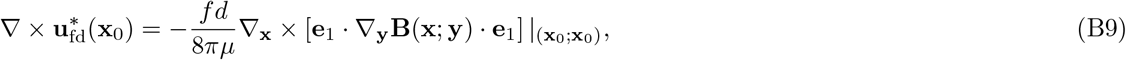

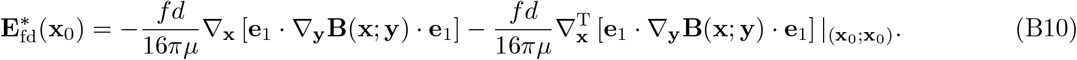

The combined effect of both singularities is obtained by adding up their respective contributions to the flow by linearity. In the numeric studies, we integrate (B1)) and (B2)) for the trajectory with the Matlab ode45 routine.

## Appendix C: Linear Stability Analysis

We aim to understand analytically the effect of the far-field singularities on the trajectory of a swimmer of typical size *L* when swimming (nearly) perpendicular or parallel to a no-slip surface. Such singularities consist of a force dipole of strength *fd*, a rotlet dipole of strength *Gw*, and terms smaller by a factor 𝒪 (*L/* ∥**x**∥), which we neglect. Despite the rotlet dipole being asymptotically smaller, it becomes comparable to the force dipole at a distance ∥**x**∥ = 𝓁 ∼*Gw/fd* from the body. This implies that for “strong feeders/expellers” with |𝓁| ≫ *L*, the rotlets govern the impact dynamics. Indeed, the singularity description is still accurate for such an organism as distances from the wall in the range *L*≪ *z* ≪ |𝓁|. Figure 5 shows that such singularities convincingly capture the flow around *C. reinhardtii*, including the stagnation point. This motivates using the above far-field description of pullers in the calculations.

**FIG. 5.**
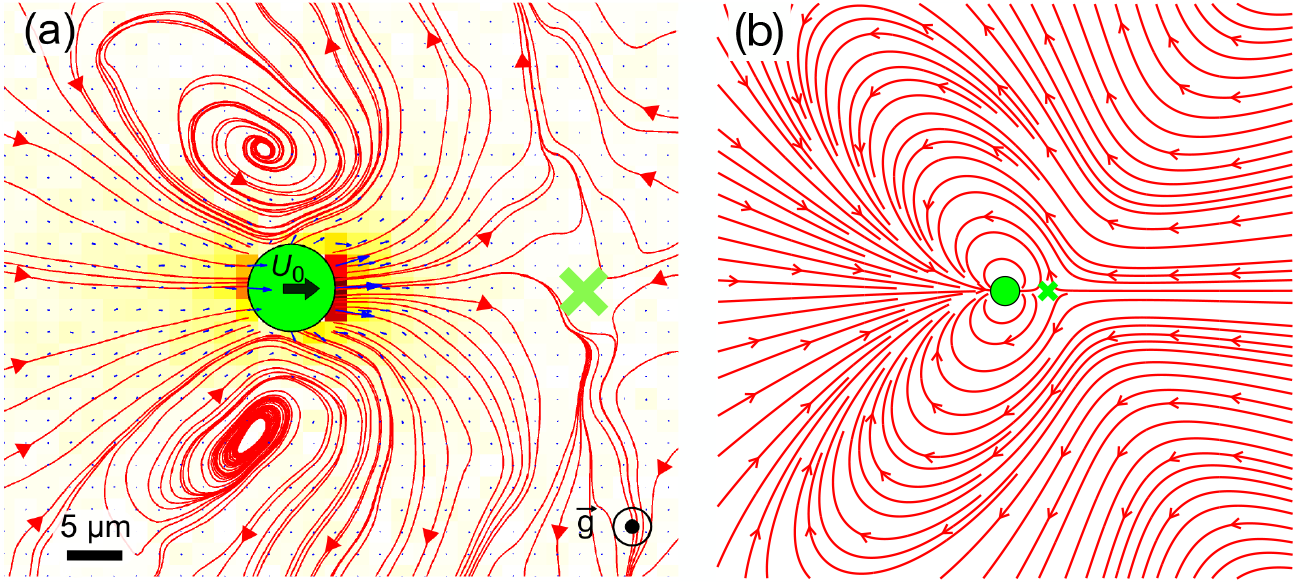
Test of the far-field model. (a) Experimental flow field around *C. reinhardtii*, from [3]. (b) Approximation consisting of two singularities located at the cell-center, a force dipole and a rotlet dipole. This differs from the approach in [3] in which three separated singularities were used.

### 1. Dynamics for a Nearly-Perpendicular Trajectory

A swimmer initially moving perpendicularly to the wall, i.e. with 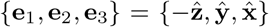, will continue to move perpendicularly by symmetry. In this section we analyze the evolution of a small perturbation

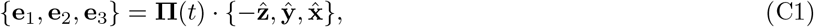

for *t* ≥ 0, where 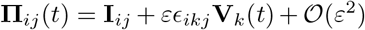, with |*ε*| ≪ 1, is an infinitesimal rotation matrix. All equations are up to 𝒪 (*ε*^2^), since {**e**_1_, **e**_2_, **e**_3_} must have unit length. From (B3) and (B7) we obtain the leading-order velocity

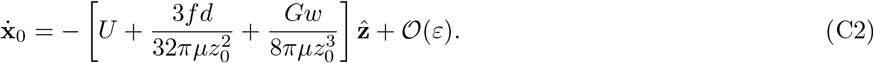

Thus, feeders (*G >* 0) receive a boost from the suction flow setup by the rotlets, while expellers (*G <* 0) are slowed down. If *G* is large and negative, eventually 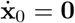 at leading order, so the organism hovers above the wall. Turning our attention to the rotational dynamics, from (B5), (B6), (B9) and (B10) the rate of turning 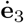 is

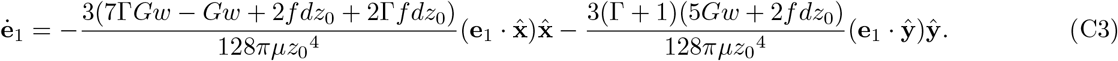

The rotifer is thus rotated towards the wall when 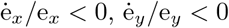, which is the case when

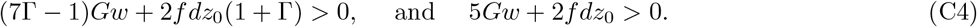

Equation (C4) shows that, unless the body is very nearly spherical, strong expellers are rotated away from the wall while strong feeders are rotated towards the wall. Rotation towards the wall is facilitated by shear alignment and impeded by the vorticity. Indeed, when 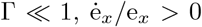 for *G* large and positive, while if Γ∼ 1 both e_*x*_ and e_*y*_ shrink over time. For rotifers, Γ∼ 0.9 *>* 1*/*7, so the rotlets have a stabilizing effect.

### 2. Dynamics for a Parallel Trajectory

Similarly to section C 1, we compute the translational and orientational dynamics for swimming parallel to the surface, i.e. when

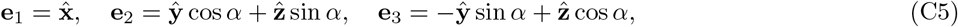

where *α* ∈ [0, *π/*2] is the “roll” angle around the body axis. Unlike in C 1, the material frame (C5) rotates even in the absence of an initial perturbation. From (B3) and (B7) we obtain the leading-order instantaneous translational velocity, while (B5), (B6), (B9) and (B10) provide the instantaneous rate of turning towards the wall

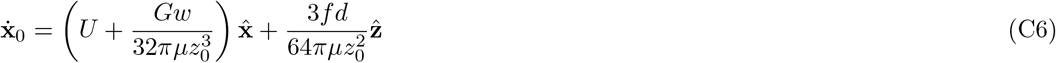

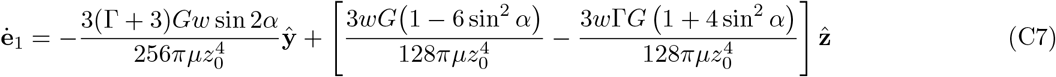

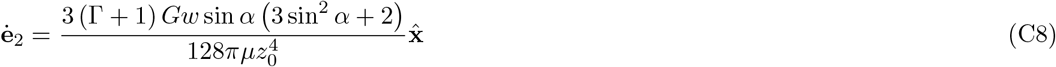

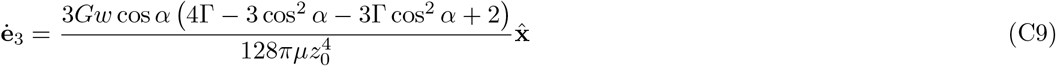

As expected, the “puller” dipole is repelled by the surface, but it is not rotated by the image flow. The rotation is instead driven entirely by the rotlet dipole flow. A feeder (*G >* 0) experiences a speed boost along the swimming direction, and is rotated towards the wall 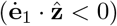 provided that

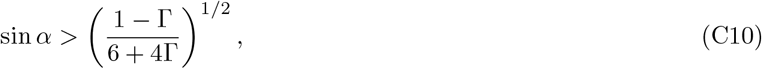

where the right-hand-side ranges from 0 (when *e* = 0) to 6^−1*/*2^ ∼0.41 (when *e* = 1). As in the orthogonal case, rotation towards the wall is promoted by shear alignment and, when sin *α* ≪ 1, is impeded by the vorticity. Therefore, for most roll angles feeders are attracted to the wall and expellers are repelled by the wall.

The hydrodynamic mechanism for reorientation is explained by examining the flow streamlines, plotted in Fig. 6 for strong expellers and feeders. Feeders facing towards the wall are sucked in, while expellers are slowed down by the outgoing rotlet flow. As for swimmers oriented parallel to the wall, the puller dipole tends to rotate the swimmer towards the wall in both case, but the rotlets aid rotation for feeders and impede it for expellers.

**FIG. 6.**
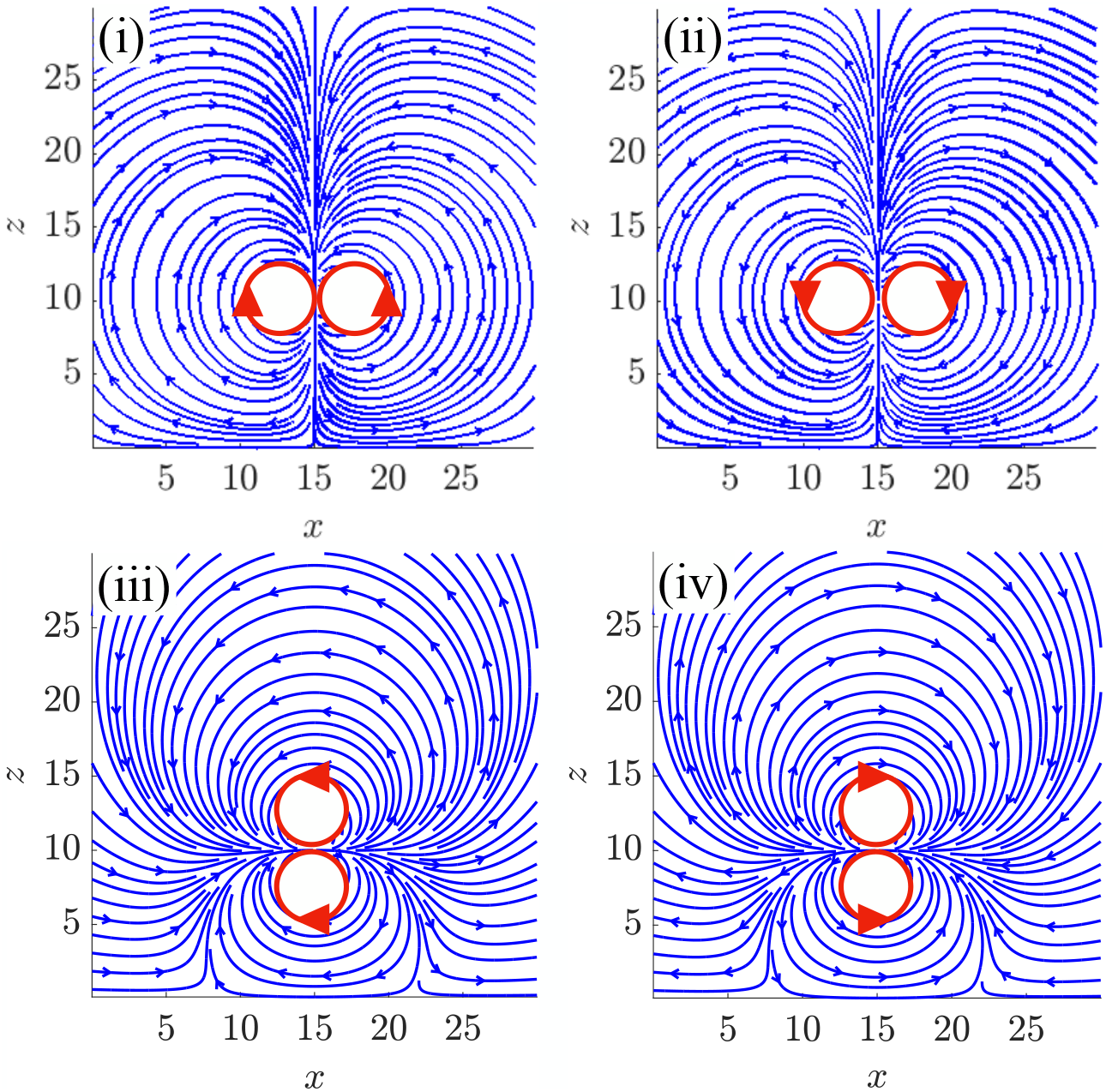
Flow produced by a rotlet dipole above a no-slip wall at *z* = 0: (i) Expeller flow when the swimmer is facing towards the wall; (ii) Feeder flow when the swimmer is facing towards the wall; (iii) Expeller flow when the swimmer is parallel to the wall; (iv) Feeder flow when the swimmer is parallel to the wall

## Appendix D: Continuum Run-and-Stick Process

In order to solve (12a) and (12b) for a function *g*(**x**, *t*) we define the Laplace-Fourier transform (LFT) 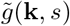 by

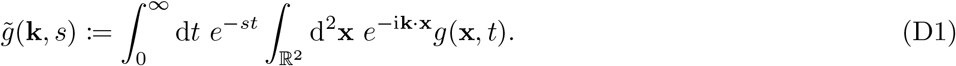

If we assume the initial conditions *f* ^+^(**x**, *θ*, 0) = δ^(2)^(**x**)*/*2*π, f* ^−^(**x**, *t*) = 0, corresponding to all rotiers released from the origin with uniformly random orientations, taking the LFT of (12a) and 12b gives

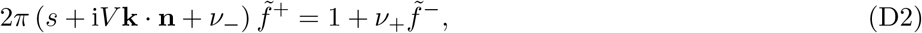

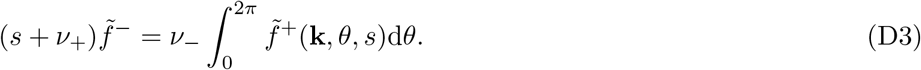

Using (D3) to eliminate 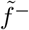 in (D2) and integrating over *θ* we obtain

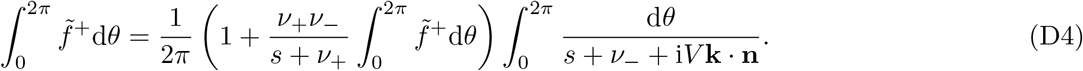

The integral may be evaluated by letting *z* = *e*^i*θ*^ and using the residue theorem, yielding

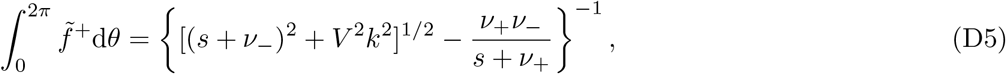

where *k* = ∥**k**∥. We may now evaluate 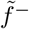 from (D3) and express the LFT of the total number density as

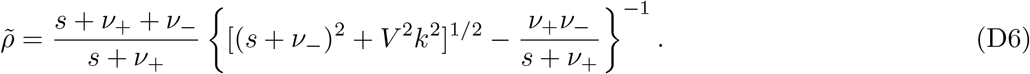

Denoting the Laplace transform by ℒ, we may exploit the rotational symmetry of *ρ* to write the MSD directly in terms of 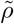,

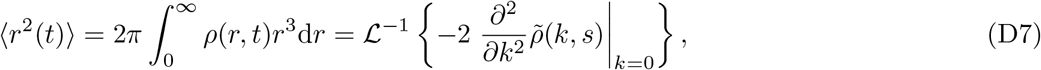

Evaluating the inverse Laplace transform by means of (D6), we obtain the analytical expression (15) for the MSD given in the main text.

